# Comprehensive and quantitative analysis of intracellular structure polarization at the apical–basal axis in elongating *Arabidopsis* zygotes

**DOI:** 10.1101/2023.08.22.554231

**Authors:** Yukiko Hiromoto, Naoki Minamino, Suzuka Kikuchi, Yusuke Kimata, Hikari Matsumoto, Sakumi Nakagawa, Minako Ueda, Takumi Higaki

**Affiliations:** Graduate School of Science and Technology, Kumamoto University, 2-39-1 Kurokami, Chuo-ku, Kumamoto, Japan; Graduate School of Life Sciences, Tohoku University, Sendai 980-8578, Japan; Suntory Rising Stars Encouragement Program in Life Sciences (SunRiSE); International Research Organization for Advanced Science and Technology, Kumamoto University, 2-39-1 Kurokami, Chuo-ku, Kumamoto, Japan

**Author notes:** Correspondence and requests for materials: Dr. Takumi Higaki Graduate School of Science and Technology, Kumamoto University, Kurokami, Chuo-ku, Kumamoto 860-8555, Japan.

## Abstract

A comprehensive and quantitative evaluation of multiple intracellular structures or proteins is a promising approach to provide a deeper understanding of and new insights into cellular polarity. In this study, we developed an image analysis pipeline to obtain intensity profiles of fluorescent probes along the apical–basal axis in elongating *Arabidopsis thaliana* zygotes based on two-photon live-cell imaging data. This technique showed the intracellular distribution of actin filaments, mitochondria, microtubules, and vacuolar membranes along the apical–basal axis in elongating zygotes from the onset of cell elongation to just before asymmetric cell division. Hierarchical cluster analysis of the quantitative data on intracellular distribution revealed that the zygote may be compartmentalized into two parts, with a boundary located 43.6% from the cell tip, immediately after fertilization. To explore the biological significance of this compartmentalization, we examined the positions of the asymmetric cell divisions from the dataset used in this distribution analysis. We found that the cell division plane was reproducibly inserted 20.5% from the cell tip. This position corresponded well with the midpoint of the compartmentalized apical region, suggesting a potential relationship between the zygote compartmentalization, which begins with cell elongation, and the position of the asymmetric cell division.

## Introduction

Cellular polarity refers to the asymmetric distribution of cellular components such as organelles and molecules. It is found in various organisms and plays essential roles in several biological processes, including morphogenesis and environmental responses^1-3^. In flowering plants, the apical–basal axis is the vertical polarity between the apical side of the shoot apex and the basal side of the root apex. The apical–basal axis is already distinct in the *Arabidopsis* zygote, with the apically positioned nucleus and basally positioned vacuole^4,5^. The zygote undergoes temporal cell disorganization just after fertilization, and then starts directional cell elongation and subsequent asymmetric cell division to produce an apical cell/proembryo and a basal cell/suspensor^6^.

Recent two-photon live-cell imaging observation demonstrated the dynamic behaviors of intracellular structures from fertilization to asymmetric cell division^7^. For example, microtubules form a subapical transverse band at the cell cortex during the cell elongation phase, and a preprophase band and phragmoplast during the subsequent cell division phase^8^. Another cytoskeleton component, actin filament, is longitudinally aligned along the apical–basal axis in the mature zygote^8^. Furthermore, the vacuole occupies a large portion of the basal side of the zygote and dynamically changes its shape and size in an actin-dependent manner^9^. Analyses using vacuolar-defective mutants revealed that the vacuole distribution plays important roles in the determination of zygote polarity^9,10^. Mitochondria also exhibited distinctive dynamics, such as filamentous elongation along the actin cables, temporal fragmentation during cell division, and uneven distribution between the apical and basal cells^11^. These spatial distributions of cellular components represent changes in the intrinsic state of the zygote during apical–basal axis formation. However, to date, the relationships among these intracellular structures (e.g., whether they are co-localized or mutually exclusive) have not been examined. Therefore, we quantitatively characterized the profiles of intracellular structure contents along the apical–basal axis to provide new insights into the polarization and compartmentalization of the *Arabidopsis* zygote.

It is challenging to simultaneously visualize multiple intracellular structures in a living cell using various fluorescent proteins and a multi-wavelength observation system. Image analysis techniques can be employed to investigate the relationships among localizations of various intracellular structures using single-channel image datasets. However, the primary challenge with current image analysis approaches is variability between samples^12^; cell size and shape can vary among individuals, and fluorescence intensity can also differ among individuals and marker probes. Therefore, normalizing cell shape and fluorescence intensity is a crucial issue for comprehensive analysis.

In a previous study, Higaki et al. (2012) collected images of 18 types of intracellular structures from *Arabidopsis* stomatal guard cells. They obtained probability maps that visualized the average distribution of these intracellular structures by normalizing both cell shape and fluorescence intensity^13^. Using subtractive images of the probability map for open and closed stomata, they visualized changes in intracellular localization during stomatal movement. This analysis identified novel relocation of the endoplasmic reticulum during stomatal opening, the dynamics of which were then experimentally confirmed^13^.

Pécot et al. (2018) presented a semiparametric framework for quantitative analysis of the distribution of fluorescently labeled proteins that considered cell shape diversity^14^. They computed 3D histograms for each parameter (radius, angle, depth) in cylindrical coordinates with cell shape normalization, which allowed comparison between cells of three different shapes^14^. Using this framework, they performed a comprehensive analysis and visualization of the spatiotemporal distribution of intracellular events in HeLa cells, and showed that cell shape and actin organization regulate Rab11 protein trafficking and vesicle distribution at the plasma membrane^14^.

Hartmann et al. (2020) developed a new point cloud-based morphometrics algorithm to transform the information in arbitrary 3D fluorescence distributions of each cell into 1D feature vectors (feature embedding)^15^. Using this algorithm, they performed a quantitative and unbiased analysis of heterogeneous cell shapes and subcellular structures in zebrafish dorsal line primordia. Furthermore, integration of embedded features using machine learning enabled an integrated analysis of cell morphology, subcellular organization, and gene expression, which is expected to contribute to the future construction of a comprehensive tissue atlas of the zebrafish dorsal line primordia^15^.

These reports underscore the feasibility and value of comprehensive image analysis techniques and image data normalization to characterize various biological problems. Consequently, we established a new comprehensive image analysis method to provide fresh insights into cellular polarity in *Arabidopsis* zygotes. In this study, we developed a new image analysis pipeline for comprehensive and quantitative assessment of the distribution of multiple intracellular structures labeled by fluorescent probes in elongating *Arabidopsis* zygotes. This pipeline quantitatively evaluates the distribution of intracellular structures along the apical–basal axis using time-sequential imaging data that account for variations in cell shape, fluorescence intensity, and probe type. Machine learning analysis of the quantitative data indicated that the *Arabidopsis* zygote was compartmentalized by the intracellular structures from the onset of cell elongation to just before asymmetric cell division. This compartmentalization may contribute to the positioning of the cell division plane during asymmetric division. This study provides new insights into plant zygote polarization through our comprehensive and quantitative image analysis technique.

## Results

### Quantitative evaluation of the intracellular structure distribution along the biological axis

From the onset of cell elongation to just before asymmetric cell division, *Arabidopsis* zygotes undergo cell elongation with nuclear migration^16^ (Fig. 1a). To evaluate the distribution of intracellular structures along the apical–basal axis during this period, we first acquired the time-lapse 3D images of actin filaments, mitochondria, microtubules, and vacuolar membranes in *Arabidopsis* zygotes from temporal cell disorganization just after fertilization to after asymmetric cell division using two-photon excitation microscopy^8,9,11^ (Fig. 1b). Then, time frames from the onset of cell elongation and just before asymmetric cell division were manually determined and trimmed. The inspection of three biological replicates for the four marker lines showed that the time duration was variable (Table 1). Therefore, we defined 10 time points by dividing this period into nine equal intervals based on two biological events: the onset of cell elongation and asymmetric cell division (Fig. 1, T = 0–9).

**Figure 1.**
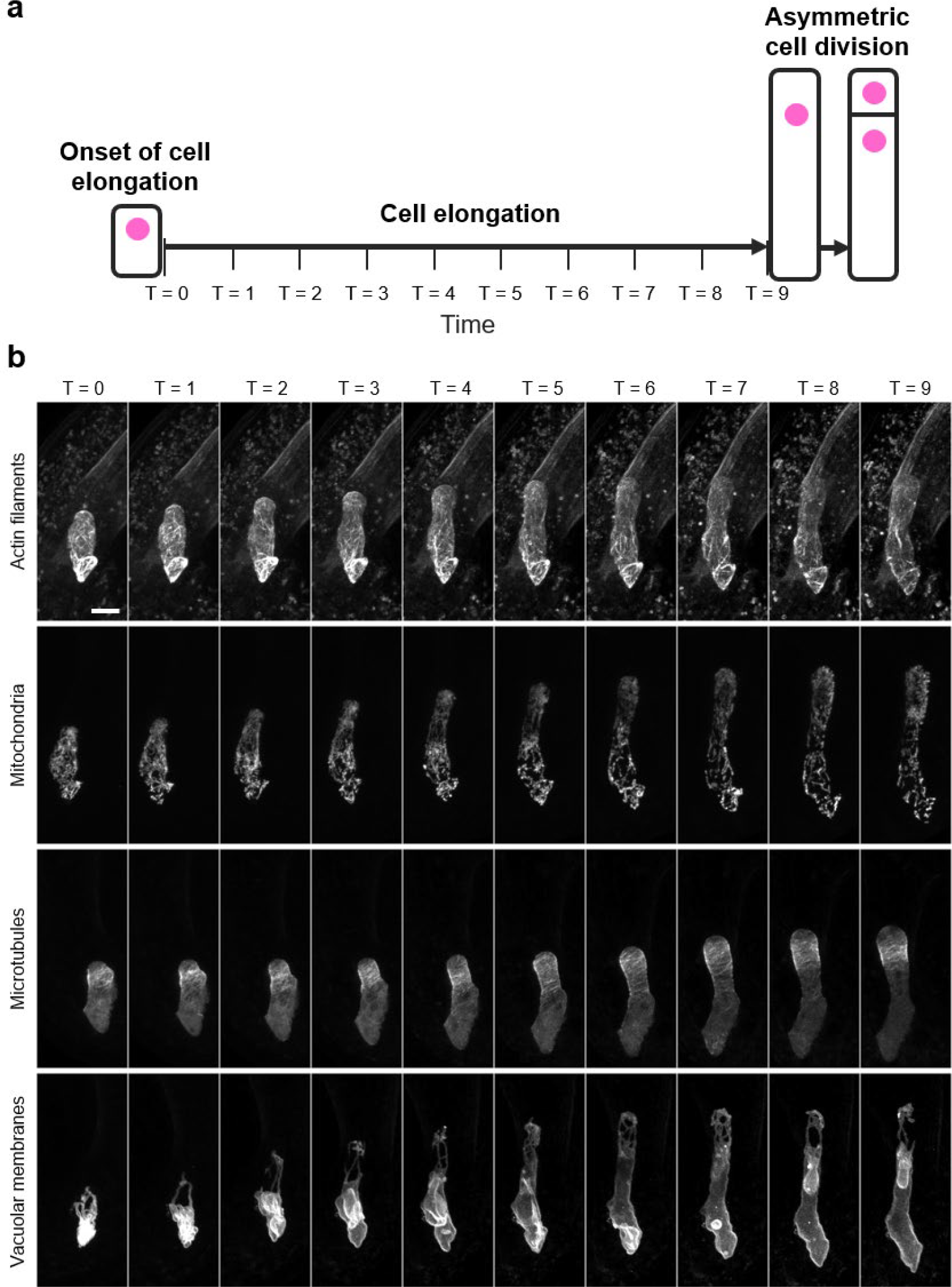
Time normalization for time-lapse images of *Arabidopsis* zygotes. (**a**) Schematic diagram of time normalization. Ten time points were defined by dividing the time period from the onset of cell elongation (T = 0) to just before the asymmetric cell division (T = 9) into nine equal intervals. Cell nuclei are illustrated in magenta. (**b**) Representative time-lapse images of time-normalized *Arabidopsis* zygotes, in which actin filaments, mitochondria, microtubules, or vacuolar membranes are fluorescently labeled. Scale bar = 10 μm.

**Table 1.** Time from the onset of cell elongation to just before asymmetric cell division in transformed zygotes used to observe an intracellular structure. n = 3. Values show mean ± standard deviation.

|  | EC1p::Lifeact-Venus for actin filaments | DD45p::mt-GFP for mitochondria | EC1p::Clover-TUA6 for microtubules | EC1p::VHP1-mGFP for vacuolar membranes |
| --- | --- | --- | --- | --- |
| Time duration (min) | 793 ± 266 | 1190 ± 61.1 | 870 ± 212 | 1290 ± 496 |

Subsequently, we extracted intensity profiles along the apical–basal axis from the 3D images. First, we acquired a maximum intensity projection (MIP) in the depth direction (Fig. 2a, b). The zygotic region was then manually segmented based on the MIP image (Fig. 2b, c). We defined the major axis of the fitted ellipse of the zygotic region as the apical–basal axis. The image was rotated so that the cell apex was oriented upwards, and the intensity of the background region was set to 0 (Fig. 2d). Following this, the average intensity profile along the apical–basal axis was obtained (Fig. 2e). Finally, intensity was normalized (mean intensity = 0, standard deviation = 1), and cell size was standardized by adjusting the cell length to a minimum value of 110 pixels (Fig. 2f). Our image processing pipeline allowed a quantitative depiction of the distribution of intracellular structures labeled with a fluorescent probe at 10 time points during zygote elongation.

**Figure 2.**
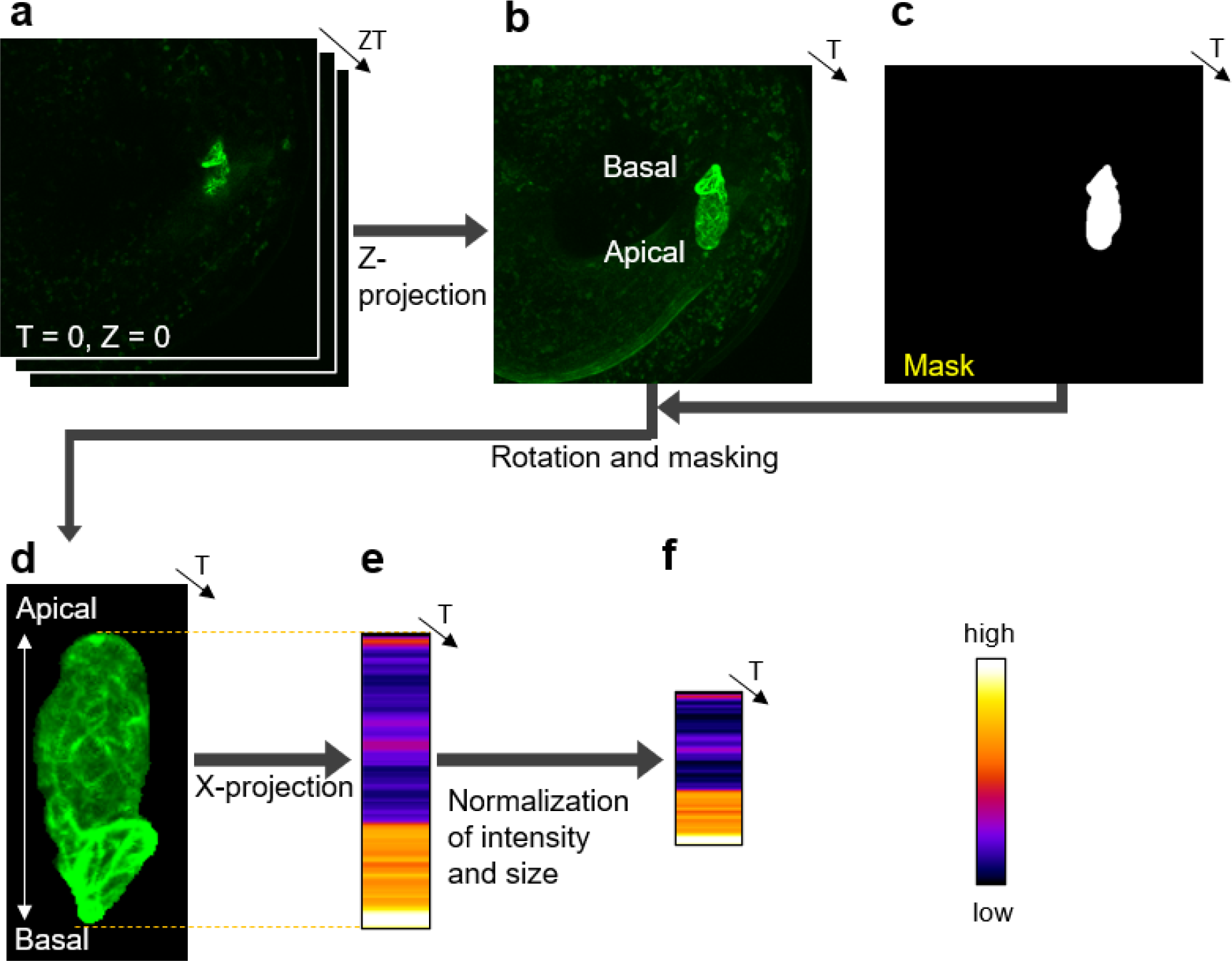
Image processing to quantitatively evaluate the distribution of the intracellular structures along the apical–basal axis. (**a**) Representative Z-stack image of the intracellular structure in an *Arabidopsis* zygote. (**b**) Maximum intensity Z-projection (MIP) image of (a). (**c**) Masked image of the zygotic region manually identified based on (b). (**d**) An image derived from the MIP image (b) wherein the zygote has been isolated from the surrounding background using the mask image (c) and rotated to align the apical–basal axis vertically with the apex oriented upwards. (**e**) Average intensity profile along the apical–basal axis. The mean intensity was calculated by averaging the intensity values of pixels located in the same position along the apical–basal axis. Note that the intensity measurement range corresponds to the cell length (broken lines). (**f**) Average intensity profile after normalization for intensity and standardization for cell size. Intensity was normalized to mean = 0 and standard deviation = 1. Cell size was standardized to 110 pixels, which was the minimum cell length examined in this study.

### Machine learning of zygote subcellular regions based on intracellular structure distribution

To visualize the changes in the intracellular structure distributions along the apical–basal axis from the onset of cell elongation to just before asymmetric cell division, we obtained a map of the intensity profiles that was organized by time points, biological replicates, and marker probes (Fig. 3a). The map showed that each intracellular structure exhibited a different distribution pattern. For example, microtubules were observed to be localized at the apical side and vacuolar membranes at the basal side (Fig. 3a). The temporal changes associated with cell elongation were not as prominent in these intracellular structures (Fig. 3a). Our findings indicate that early establishment of an uneven distribution pattern of the intracellular structures in the zygote exists when the zygote starts elongation; this pattern is maintained until just prior to asymmetric cell division.

**Figure 3.**
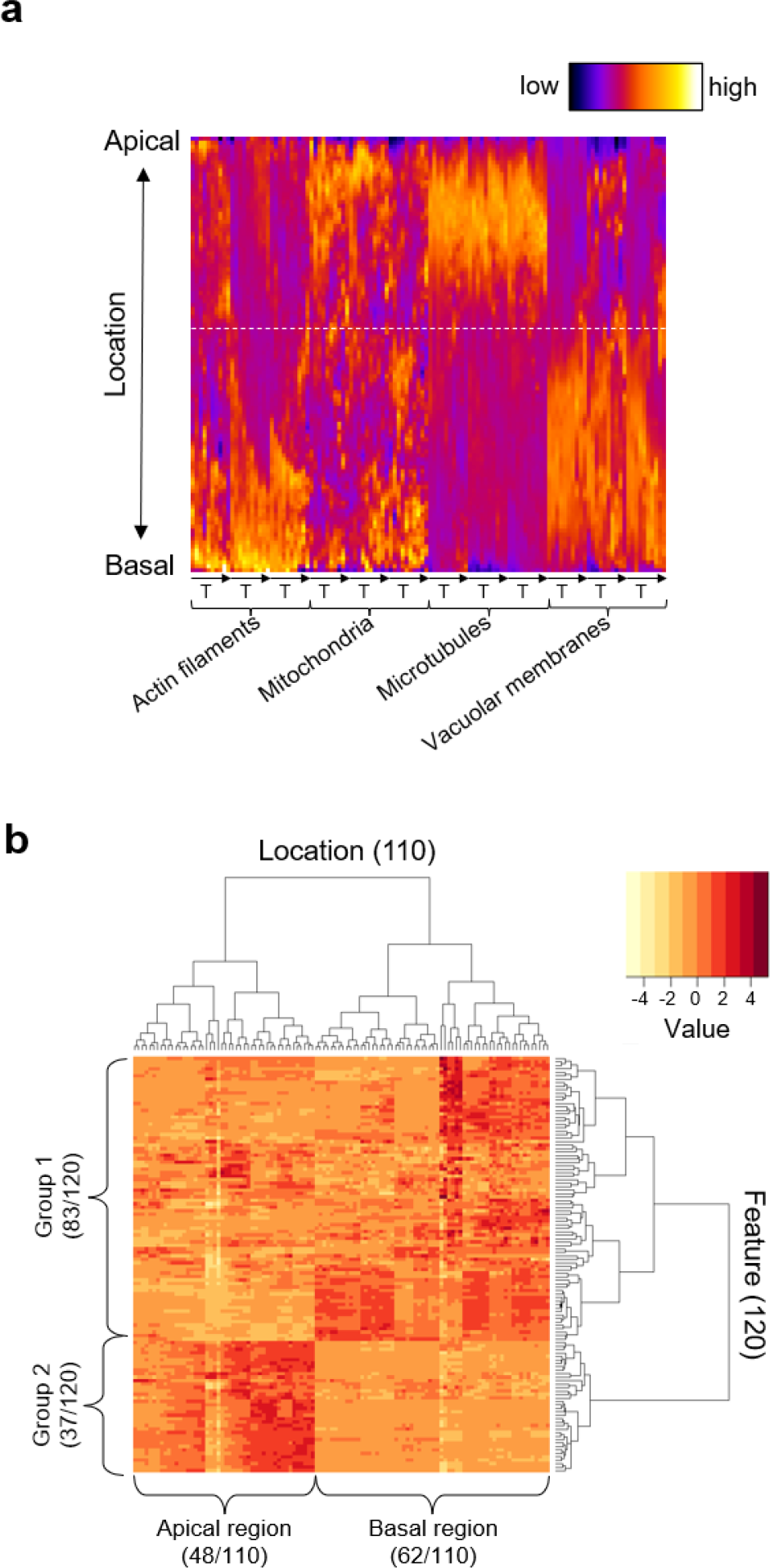
Compartmentalization of zygotes by the distribution of intracellular structures. (**a**) Spatiotemporal distribution maps of intracellular structures represented by normalized and standardized average intensity profiles. The vertical axis represents the locations along the apical– basal axis of the zygotes (110 pixels/points from the base to the tip). The horizontal axis represents 120 features, which correspond to ten time points, three biological replicates, and four marker probes. The broken white line indicates the boundary between apical and basal subregions identified by hierarchical clustering. (**b**) Dendrogram with a heat map obtained by hierarchical clustering. The top part of the dendrogram represents the clustering results of intracellular locations (Location; 110 points), and the right part of the dendrogram represents the clustering results of the probe intensities (Feature; 120 intensities).

To explore the potential compartmentalization of the zygote by the distribution of intracellular structures, we examined the positions along the apical–basal axis of the zygote—110 points from the base to the tip. These positions were characterized by the distribution of four intracellular structures at 10 time points, with three biological replicates (i.e., 120 features), and their similarity was assessed using hierarchical clustering, which is a kind of unsupervised learning^17^ (Fig. 3b). We further analyzed the categorization of each point into two clusters. The results indicated that the 110 points from the tip to the base bifurcated around the 48th and 49th points, suggesting that the zygote may be compartmentalized approximately 43.6% (48th point/110 points) from the tip (Fig. 3a, white broken line; Fig. 3b, apical and basal regions). In addition, through hierarchical clustering, intensities of the fluorescent probes labeling the intracellular structures at each time frame—used as features for identifying the compartmentalization—could be classified into two groups (Fig. 3b, Groups 1 and 2). Groups 1 and 2 exhibited higher values in the basal and apical regions, respectively (Fig. 3b).

To verify the accumulation of intracellular structures in the apical and basal regions and assess their temporal dependence, we also examined the composition of these features (Figure 4). We found that all actin filaments and vacuolar membranes belonged to Group 1, whereas microtubules were exclusively in Group 2 (Fig. 4a). We did not observe any specific time frames (T = 0–9) that predominantly clustered in either Group 1 or 2 (Fig. 4b). These results indicate that, from the onset of cell elongation, zygotes become polarized, with microtubules localized at the tip and actin filaments and vacuoles at the base, and this polarity is maintained until just before asymmetric cell division occurs.

**Figure 4.**
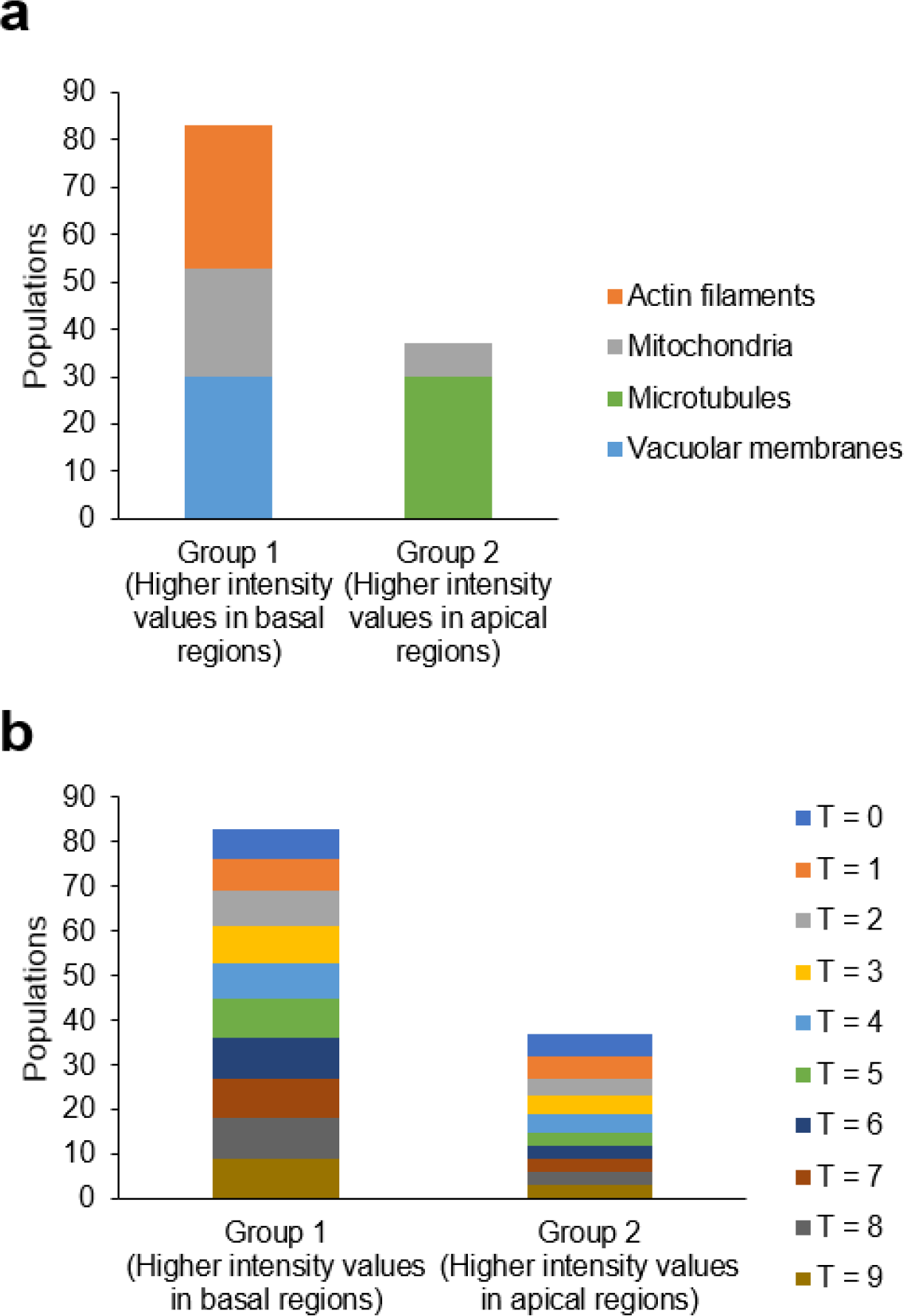
Composition analyses of features used to examine the compartmentalization of *Arabidopsis* zygotes. (**a**) Composition of intracellular structures in Groups 1 and 2, which showed higher intensity values in the basal and apical regions, respectively. (**b**) Composition of time frames (T = 0–9) in Groups 1 and 2.

To determine which of the four types of intracellular structures contribute to the compartmentalization of the zygotes, we conducted a supervised learning analysis. A classification model to segregate the apical and basal subregions of the zygote was developed using the “random forest” method, an ensemble machine learning algorithm that enhances generalization capabilities by integrating multiple weak decision tree learners^18,19^. The classification accuracy was evaluated using the out-of-bag error rate, which is computed by classifying the training data as test data subsequent to the construction of a partial forest by aggregating groups of decision trees not involving the training data. The out-of-bag error rate was 1.8% (two errors out of 110 points) (Table 2), which indicated high classification accuracy.

**Table 2.** Confusion matrix of the *Arabidopsis* zygote apical and basal subregion supervised classification. Out of the 110 points examined, only two were incorrectly classified; this resulted in an error rate of 1.8%.

|  | Predicted apical region | Predicted basal region |
| --- | --- | --- |
| Apical region | 47 | 1 |
| Basal region | 1 | 61 |

Additionally, the random forest algorithm can estimate the importance of the variables used for classification^18^. Consequently, we investigated the relative importance of these four intracellular structures based on the mean decrease in the Gini coefficient^19^. Vacuolar membranes were identified as the most significant determinant for the classification of apical and basal regions (Fig. 5a). Moreover, no significant difference was detected in the time frame contribution to the classification (T = 0–9) (Fig. 5b).

**Figure 5.**
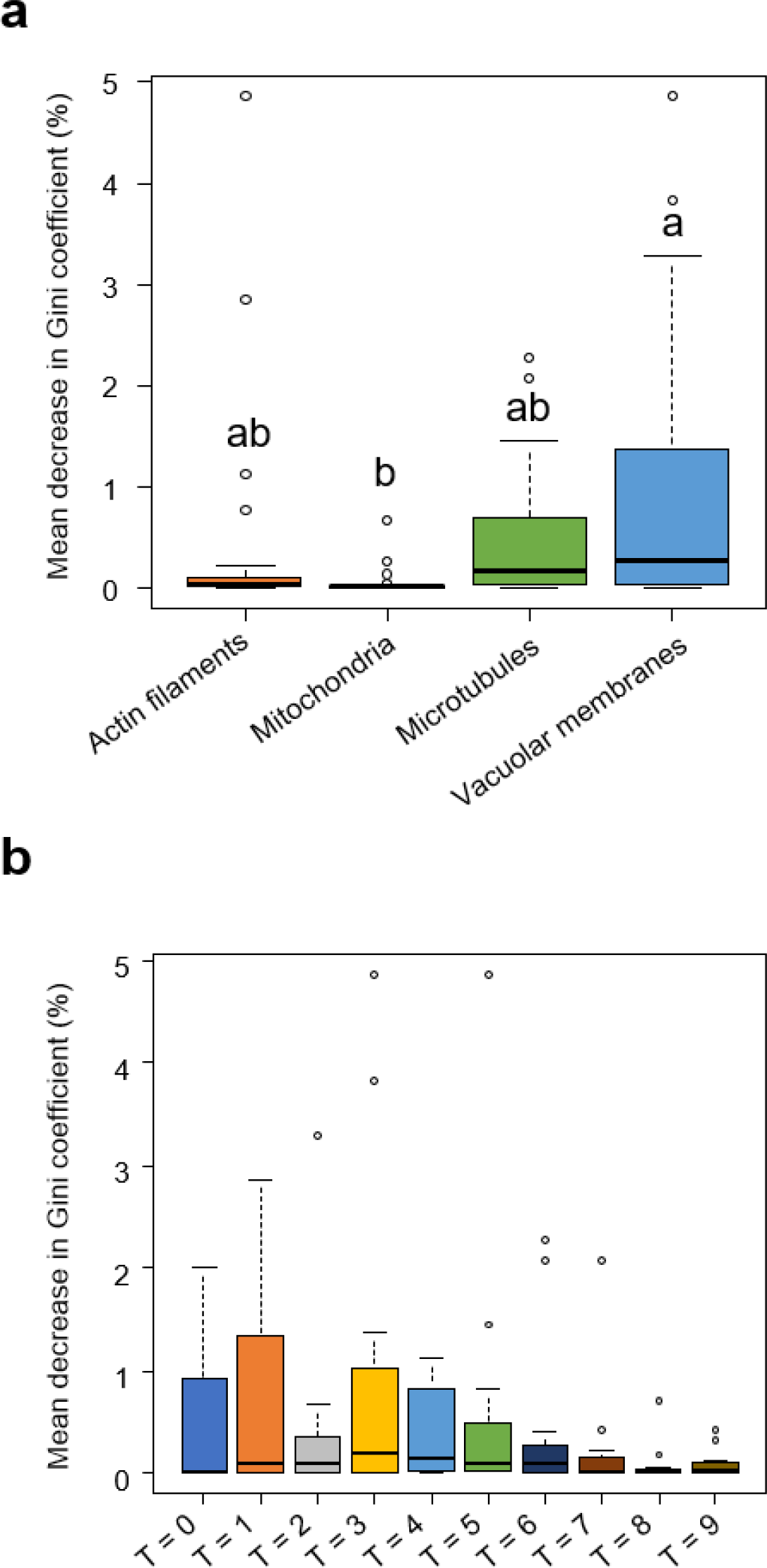
Relative importance of features for segregating the apical and basal regions in elongating *Arabidopsis* zygotes using a random forest classification model. **(a, b)** Relative importance of individual intracellular structures (**a**) and time frames (**b**) for the classification for the apical and basal regions. Box plot showing the mean decrease in Gini coefficient (%) for each intracellular structure. A higher value of the mean decrease in Gini coefficient indicates more substantial contribution to random forest classification. Different alphabets indicate significant differences (n = 3. Tukey–Kramer test, p < 0.01).

### Correlation between zygote compartmentalization and asymmetric cell division planes

To elucidate the biological significance of the *Arabidopsis* zygote compartmentalization, we explored its correlation with the position of the asymmetric cell division plane. We manually identified the location of the asymmetric cell division plane from our time-lapse data set, and quantified its relative position on the apical–basal axis (Fig. 6a). Our findings demonstrated that the division plane was inserted at an average position of 20.5% ± 2.15% (mean ± SD) from the cell tip (Fig. 6b). This position aligns with half of the compartmentalized apical region (43.6% from the tip), which indicates a potential relationship between zygote compartmentalization and the positioning of the asymmetric cell division plane (Fig. 7).

**Figure 6.**
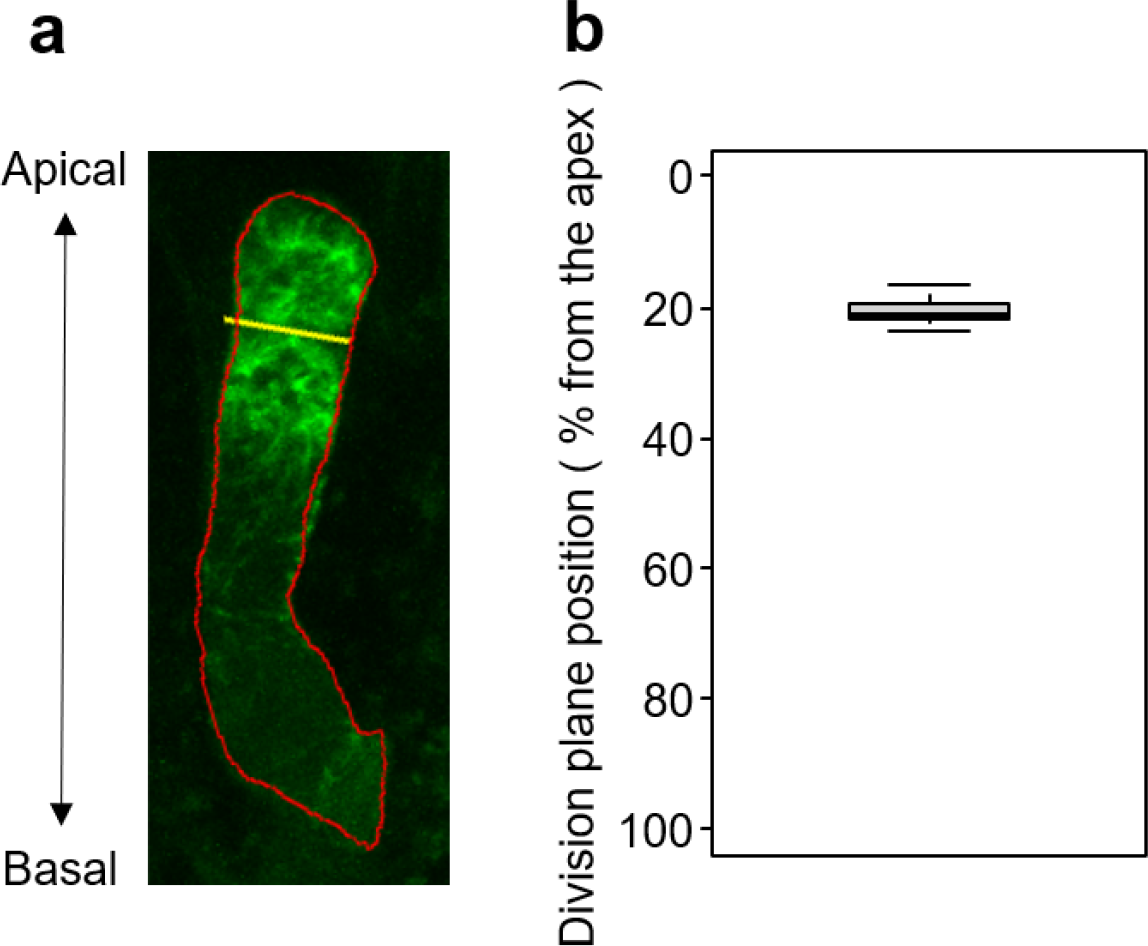
Position of the asymmetric cell division plane. (**a**) Representative image of a zygote expressing the microtubule marker Clover-TUA6 (green) with a defined asymmetric cell division plane. Cell designation (red) and a division plane (yellow) were manually determined. (**b**) Box plot showing the position (% from the tip) of an asymmetric cell division plane. n = 12. Mean ± standard deviation = 20.5% ± 2.15%.

**Figure 7.**
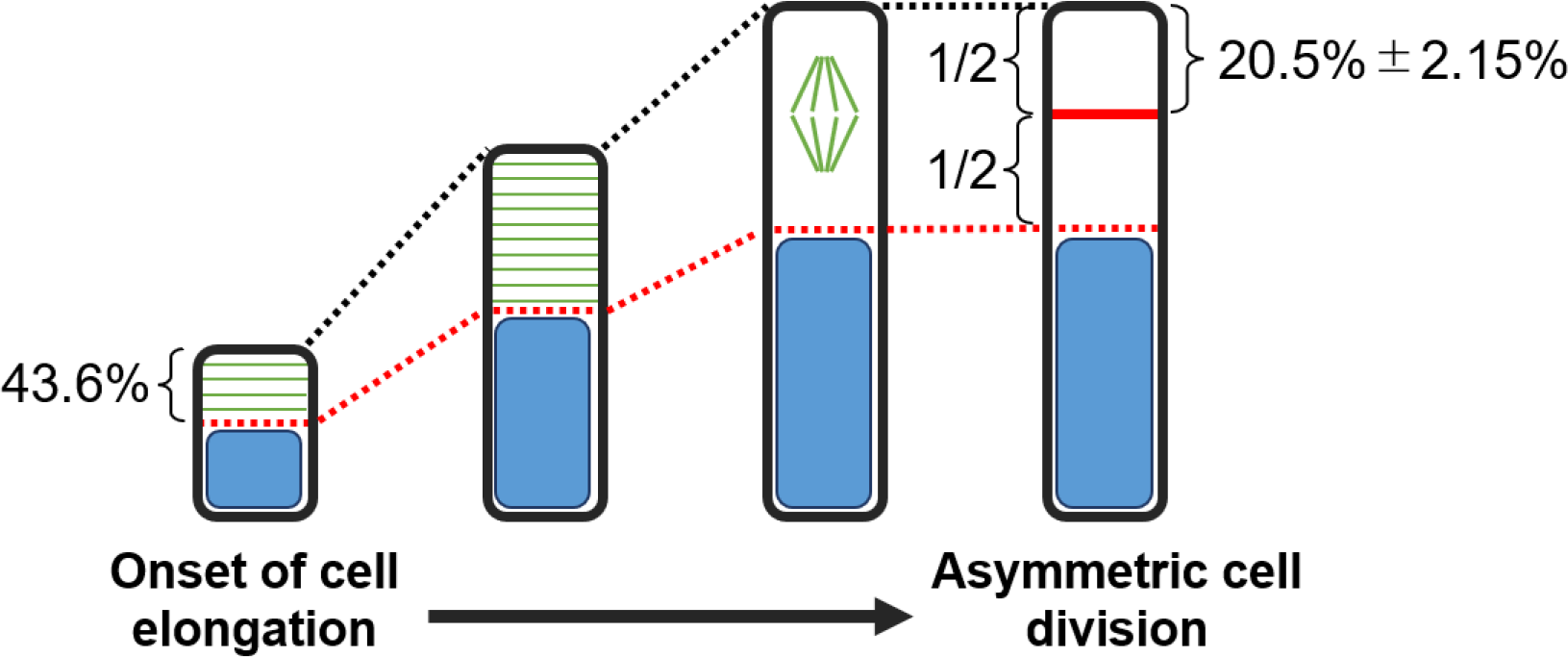
Hypothetical schematic illustration of polarization in *Arabidopsis* zygotes from the onset of cell elongation to asymmetric cell division. For clarity, only vacuoles (blue) and microtubules (green), which indicate polarization, are shown here. At the onset of cell elongation, the zygote is already compartmentalized 43.6% from the tip, and the proportions of this compartment are maintained through asymmetric cell division (red broken line). A correlation was observed between the position of the apical compartment’s midpoint and the division plane during asymmetric cell division (red solid line).

## Discussion

In this study, we quantitatively evaluated the polarized localization of the intracellular structures using an in-house-developed image analysis pipeline. This pipeline can comprehensively and quantitatively analyze the time-lapse 3D images of multiple *Arabidopsis* zygotes exhibiting diverse time scales, fluorescence intensity, and cell sizes (Fig. 2). Our image analysis pipeline allowed us to visualize and quantitatively describe the distribution of multiple intracellular structures along the apical–basal axis in the *Arabidopsis* zygote (Fig. 3a). The clustering analysis based on the obtained distribution information revealed that *Arabidopsis* zygotes may be compartmentalized from the onset of cell elongation to just before asymmetric cell division, featuring an enrichment of microtubules in apical regions and actin filaments and vacuoles in basal regions (Figs. 3 and 4). Our findings are consistent with previous observations of individual probes, such as a cortical microtubule band at the apical region^8^, large vacuoles, and actin filaments that regulate vacuolar shape at the basal region^9,10^, and a dispersed distribution of mitochondria^11^.

We constructed a comprehensive view of zygote polarization orchestrated by these intracellular structures. Our method requires accurate segmentation of cellular regions. In this study, manual segmentation was performed because of its importance in ensuring segmentation accuracy. However, depending on the volume of data, the introduction of an automatic segmentation method^16^ may be useful. A more fundamental limitation of our method is its inability to provide information beyond two dimensions because of the dimensionality reduction that occurs during the image processing phase, which results in one-dimensional data. Although this study focused on the apical–basal axis, a different approach would be required if two or more biological axes were simultaneously considered. Nevertheless, our pipeline is distinguished by its simplicity and versatility; in particular, it requires no complex algorithms or parameter inputs, especially compared with the comprehensive analysis methods proposed in previous studies^14,15,20^. Therefore, our pipeline is helpful because it is universally applicable to *Arabidopsis* zygotes in addition to other species and cell types that exhibit a biological axis. For example, at the subcellular level, it may be useful to analyze protein dynamics during asymmetric cell division, which occurs during *Bacillus subtilis* endospore formation^21^ and *Caenorhabditis elegans* embryo development^22^. It is also possible to perform analyses at scales higher than the cellular level. For example, this approach could contribute to a comprehensive analysis of the spatiotemporal dynamics of phytohormone responses in the apical–basal axis of the root at an organ level.

Our hierarchical clustering analysis using the spatiotemporal information derived from subcellular distribution of each intracellular structure showed that the zygote could be compartmentalized with a boundary 43.6% from the tip of the apical end (Fig. 3). Notably, this compartmentalization formed early in the process and was maintained until just before asymmetric cell division. Moreover, despite the zygote’s dramatic elongation during this period, the proportion of intracellular compartmentalization remained constant (Fig. 7). These findings demonstrate the potential importance of early zygotic polarization in plant development. This notion is supported by a previous finding that mutants displaying more aberrant asymmetric cell division exhibit abnormalities in cell elongation, rather than in the asymmetric cell division itself^23^.

The importance analysis from random forest classification indicated that the distribution of the vacuolar membrane contributed the most to this compartmentalization among the intracellular structures (Fig. 5). Zygotic vacuoles undergo dynamic reorganization from fertilization to asymmetric cell division. Vacuoles form tubular structures around the migrating nucleus toward the apical region and become small in the apical region, whereas vacuolar size in the basal region is increased^9^. *sgr2-1* and *vti11*, which were previously reported to be defective in vacuolar function^24-27^, fail to form dynamic tubular networks of the vacuoles and exhibit abnormally large spherical vacuoles in the apical area; this leads to irregular asymmetric division and morphological abnormalities in the zygote^9,10^. These observations indicate that appropriate vacuolar distribution plays a crucial role in the proper development of the zygote, which is consistent with our results in this study. We also noticed that the location of the asymmetric cell division plane in the zygote coincides well with the midpoint of the apical compartment region (Fig. 6). This correlation suggests that the polarized distribution of intracellular structures may be associated with determining the site of asymmetric cell division during zygotic development (Fig. 7). At present, a correlation has only been found between the compartmentalization and asymmetric cell division planes; therefore, we should be cautious about inferring a causal relationship between them. Nevertheless, investigating whether our method can predict the site of the division plane in mutants exhibiting aberrant asymmetric cell division, such as *ssp*^28^, *yda*^*29*^, *wrky2*^23^, and *hdg11/12*^23^, represents an interesting topic for a future study.

One notable question is whether the vacuolar distribution significantly contributes to other polar patterning resulting from asymmetric cell division. In stomatal development, asymmetric cell division of a meristemoid mother cell gives rise to a small meristemoid, which finally differentiates into stomatal guard cells through amplified divisions and differentiation steps^30-32^. Although the vacuolar-defective mutant *sgr2-1* does not exhibit any defects in stomatal development and patterning^9^, precise vacuolar contribution to asymmetric cell division in the meristemoid mother cell has yet to be ascertained. The tip-growing cells of protonemata in moss also undergo asymmetric cell division, which generates an apical cell with continual tip growth and a subapical cell containing a large vacuole^33,34^. The large vacuole is disproportionately located on the basal side, but the significance of vacuolar location for asymmetric division remains unknown^34^. Further examination of vacuolar dynamics in stomatal development in *Arabidopsis* and protonemal tip growth in moss could provide valuable insights into the determination of the asymmetric cell division plane.

## Methods

### Plant growth conditions and transgenic lines

All *Arabidopsis* lines were in Col-0 background. Plants were grown at 18–22 °C under continuous light or long-day conditions (16-hour light/8-hour dark). The actin filament marker was EGG CELL1 (EC1)p::Lifeact-Venus^35^, and the microtubule marker was EC1p::Clover-TUBULIN ALPHA6 (TUA6)^8^. The mitochondrial marker (DD45p::mt-GFP)^36^ was previously described^11^. The vacuolar membrane marker (EC1p::VACUOLAR H^+^-PPASE (VHP1)-mGFP) was previously described^9^.

### Time-lapse observation

The zygote live-cell imaging was performed using a laser-scanning inverted microscope (A1MP; Nikon) equipped with a Ti:sapphire femtosecond pulse laser (Mai Tai DeepSee; Spectra-Physics) as previously described^37,38^. Time-lapse images were taken every 20 min with 3× zoom and 31 Z-stacks with 1 μm intervals.

### Image processing

We used the time-lapse 3D (XYZ) confocal images of zygotes, in which actin filaments, mitochondria, microtubules, and vacuolar membranes were fluorescently labeled (Fig. 1b). To examine the changes in the distribution of intracellular structures along the apical–basal axis from the onset of cell elongation to just before the asymmetric division, we manually identified the time frames of this period from the raw 4D images and then extracted 10 time frames at equal time intervals. The time frames were defined by the biological events; therefore, each cell has different time intervals. The MIP image for the Z-axis direction in each time frame was obtained (Fig. 2b). For the manual segmentation of the zygote regions, the MIP images were used (Fig. 2c).

To define the apical–basal axis, the zygote regions were fitted into ellipses. The major axis of the fitted ellipse was defined as the apical–basal axis. All projection images were masked and rotated so that the apical–basal axis was aligned with the Y-axis of the image with the cell apex positioned upward (Fig. 2, Rotation and masking).

To evaluate the intensity distribution along the apical–basal axis, the zygote pixels were averaged with respect to the axis orthogonal to the apical–basal axis (i.e., the X-axis of the image) (Fig. 2, X-projection). In addition, the length of the zygotes was normalized by setting them to 110 pixels, which was the minimum length in the zygotes observed in this study. The intensities were also normalized to an average intensity of 0 with a standard derivation of 1 (Fig. 2, Normalization of intensity and size). This image processing generated 110 positional data points along the apical–basal axis, which included 120 features consisting of four cellular structures × 10 time points × three biological replicates. Of the three biological replicates in the four probes, one replicate was identical to a replicate used in our previous papers^8,9,11^. All image processing was performed using ImageJ software version 1.53e^39^.

### Hierarchical clustering and random forest classification

Hierarchical clustering was performed to group 110 positional data points based on 120 features using R software version 4.0.5^19^. The intensity profiles were normalized using the “scale” function with a mean of 0 and a standard deviation of 1. Hierarchical clustering and acquisition of a heatmap were performed using the “heatmap.2” function with the ward.D2 method. When divided into two clusters, the assignment of each intracellular position or feature to a cluster was determined using the “cutree” function.

The model for classifying tip or base labels based on the result of the hierarchical clustering was trained using the 110 positional data points, which included 120 features, with the R package “randomForest”^19,40^ (https://cran.r-project.org/web/packages/randomForest/index.html). Two specifications were set: “Number of trees = 500” and “No. of variables tried at each split = 10.” The remaining parameters were used with the default settings. Therefore, the training data were randomly extracted from a portion of the 110 positional data points, and 10 variables were randomly selected from 120 features to generate a decision tree. This iterative process was repeated until a total of 500 decision trees were generated. The final classification result was determined by majority voting among the individual classification results. Importance was evaluated using the “importance” function as the average decrease in the Gini coefficient, and the average of the importance was measured in each cellular structure (actin filaments, mitochondria, microtubules, and vacuolar membranes).

## Data availability

Data are available upon reasonable request to the corresponding author (T. H.).

## Acknowledgments

We thank Prof. Koichi Fujimoto of Hiroshima University and Dr. Satoru Tsugawa of Akita Prefectural University for helpful discussions. We also thank Mallory Eckstut, PhD, from Edanz (https://jp.edanz.com/ac) for editing a draft of this manuscript. This work was supported by the Japan Advanced Plant Science Network, the Japan Society for the Promotion of Science [a Grant-in-Aid for Research Activity Start-up (JP21K20650 to H.M.), a Grant-in-Aid for Early-Career Scientists (JP22K15135 to H.M., and JP23K14204 to Y.K.), a Grant-in-Aid for Scientific Research on Innovative Areas (JP19H05670 and JP19H05676 to M.U.; JP16H06280 (Advanced Bioimaging Support)), a Grant-in-Aid for Scientific Research (B) (JP19H03243 to M.U.)], and the Japan Science and Technology Agency [CREST (JPMJCR2121 to T. H. and M.U.)].

## Author contributions

T.H. conceived and designed the experiments. Y.K., H.M., S.N., and M.U. acquired the microscopy images. Y.H. and T.H. performed the image analysis experiments. T.H., N.M., S.K., and Y.H. wrote the manuscript, and all authors participated in the discussion and review of the manuscript.

